# Toxin-glycolipid interactions measured by imaging surface plasmon resonance on artificial membranes predicts diffusion behavior and lipid dependence of binding to cells

**DOI:** 10.1101/2024.03.22.586282

**Authors:** Sarah Lehnert, Umit Hakan Yildiz, Natalie Haustein, Enlin Li, Artur Matysik, Kamila Oglęcka, Rafi Rashid, Elke Boschke, Bo Liedberg, Thorsten Wohland, Rachel Kraut

## Abstract

Membrane-protein interactions mediate cellular invasion by toxins, and are thought to involve organized plasma membrane lipid domains, often containing glycolipids, other sphingolipids, and/or cholesterol. Here, we characterize an isolated glycolipid-interacting domain of the tetanus toxin heavy chain (Hc) as a fluorescently labelled peptide, TeNT46, and describe its membrane dynamics and binding characteristics on artificial bilayers and cellular membranes. We show that this novel ganglioside-interacting probe TeNT46 retains the glycolipid binding preferences of the parent toxin, using imaging-SPR (iSPR) on a micro-patterned hybrid bilayer surface. On live cell membranes, using fluorescence correlation spectroscopic (FCS) diffusion measurements to compare TeNT46 to the well-studied GM1-binding toxin CTxB, we find that both probes display ordered domain-binding characteristics, but distinct cholesterol and sphingolipid dependencies. Strikingly, the contrasting lipid requirements of TeNT46 from those of CTxB in cells are predicted by their iSPR binding preferences on hybrid synthetic membranes. Based on the combined findings from iSPR and FCS, we propose a model for toxin-membrane interaction whereby a unique lipid constellation determines optimum binding for each probe independently of lateral confinement, which is more generally imposed by cholesterol. Our resulting understanding of the specific lipid requirements of these toxin targets and their dynamics in cell membranes could be important for the future design of preventive membrane-based nano-decoys and cell-delivery tools.

## Introduction

Many examples of toxin invasion into cells depend on the presence of membrane domains that contain a specific constellation of lipids, usually including sphingolipids and/or glycolipids and cholesterol (Aigal et al., 2015; Fantini, 2007; Gruenberg and van der Goot, 2006). The organization and dynamics of these membrane domains responsible for toxin binding and uptake are little understood, due to a paucity of probes by which they can be examined (Engberg et al., 2016; Kraut et al., 2012; Mizuno et al., 2011; Skocaj et al., 2013). Here, we describe a novel ganglioside-binding, fluorescently tagged peptide probe derived from a fragment of the tetanus neurotoxin Hc terminal domain. Tetanus toxin binding to gangliosides, or sialylated glycosphingolipids, via the Hc domain has been well characterized at the molecular and structural level (Chen et al., 2009, 2009; Emsley et al., 2000; Fotinou et al., 2001; Louch et al., 2002; Rummel et al., 2003; Shapiro et al., 1997), and the toxin is internalized by a sequential clathrin-dependent mechanism initiated within lipid microdomains (Deinhardt et al., 2006).

We monitored the binding of this tetanus-derived probe, TeNT46, to artificial ganglioside-containing supported bilayer surfaces of different lipid compositions, using imaging-surface plasmon resonance (iSPR). Ellipsometric imaging combined with SPR (or ‘imaging SPR’; iSPR), like its forerunner SPR, is a surface-sensing technique that detects binding of an analyte to a surface, based on changes in refractive index and the resulting changes in angle of reflection of incident light upon the surface (Howland et al., 2007). Additionally, iSPR employs ellipsometry to detect and image angstrom-range height features in the surface. This enables verification of the planar morphology of the artificial surface, an important determinant of reproducibility of the peptide-membrane interaction, and of biological verisimilitude with regard to a planar plasma membrane (Groves and Boxer, 2002). Here, we improved upon this method by building a hybrid bilayer on top of a thiol-derivatized alkane stamped in a micropattern, which enabled multiple measurements on a single chip (see (Plant, 1999) for a review of such bilayers). Each measurement could thus be normalized to a self-contained internal control, i.e. the non-stamped area.

Centrally to this study, the fluorescent tagging of TeNT46 allowed us to monitor using fluorescence correlation spectroscopy (FCS) the dynamic behaviour of the probe on live cell membranes as an independent confirmation of the iSPR hybrid bilayer binding results. Previously, we and others have used FCS on other toxin-derived, microdomain-interacting peptide probes (Bacia et al., 2004; Hebbar et al., 2008; Lauterbach et al., 2012; Sankaran et al., 2009; Steinert et al., 2008) to define a characteristic bimodal, confined-diffusion behaviour, exhibiting a slow component dependent on sphingolipids and cholesterol, as well as more complex anomalous behaviours (Day and Kenworthy, 2012; Kang et al., 2019; Lenne et al., 2006; Wawrezinieck et al., 2005). Using FCS to detect changes in the relative amount of the slower diffusing, bound fraction vs. the freely diffusing, unbound fraction of the probe molecules measured, we were also able to compare the lipid binding requirements on cells, after depletion by different pharmacological agents. These data could then be compared to the binding data derived from iSPR on synthetic membranes. Diffusion and binding behaviours exhibited by both the novel probe TeNT46 and the well-characterized CTxB on the plasma membrane of living cells bore out the lipid requirements that were suggested by iSPR binding experiments on artificial hybrid bilayers.

Using this combination of iSPR with artificial bilayers of varying compositions, and FCS on live cells before and after lipid perturbations, we discovered that the tetanus-derived probe, TeNT46, preferred a different ganglioside binding target from CTxB, for which the iSPR results identified the expected target lipid, GM1. iSPR measurements confirmed GT1b among a variety of gangliosides as the favoured binding partner for TeNT46, identical to the most frequently cited cellular receptor for the intact tetanus toxin (Fotinou et al., 2001; Louch et al., 2002; Rummel et al., 2003). Moreover, TeNT46 showed a different lipid constellation preference from CTxB required for optimal binding to the artificial membrane, in parallel with diffusion characteristics that were distinct from CTxB on the cell plasma membrane. However, our spectroscopic measurements provide evidence that both the tetanus and cholera toxin-derived probes reside in membrane domains that exhibit constrained diffusion that is dependent upon sphingolipids and/or cholesterol.

Thus, we were able to detect a surprising retention by TeNT46 of the parent toxin molecule’s binding specificity (with a preference of GT1b over GD1b and several other gangliosides) and moreover, a correlation of the lipid requirements in the artificial bilayer to those in actual cells, demonstrating the predictive power of the iSPR experiments to the diffusion behaviour of the different toxin-derived probes.

## Materials and Methods

### Lipids and peptide probes

1-palmitoyl-2-oleoyl-sn-glycero-3-phosphocholine (POPC), sphingomyelin (SM) and the gangliosides GM1, GD1a, GD1b, GT1b and GQ1b were purchased from Avanti polar lipids Inc. and Carbosynth Limited, respectively. Cholesterol (C) and 98% 1-octadecanethiol (ODT) were purchased from Sigma Aldrich. Rhodamine labeled with 1,2-dihexadecanoyl-sn-glycero-3-phosphoethanolamine (rhodamine-DPPE) was purchased from Molecular probes. The reference protein Cholera toxin subunit B (CTxB) conjugated with Alexa fluor 488 (recombinant), Hank’s balanced salt solution (HBSS) and HEPES buffer were purchased from Life Technologies. TeNT46 was synthesized by GL Biochem Ltd. (Shanghai). All chemicals were used without further purification.

### Imaging Surface Plasmon Resonance

An imaging null-ellipsometer (EP3, Nanofilm, Germany) equipped with an SPR cell was used to analyse surface patterning and hybrid bilayer formation and subsequent peptide binding to the substrate. The Kretschmann setup was used, where a 60° BK-7 prism is attached to an approximately 100 μL flow cell. The light source was provided by a xenon lamp, with a wavelength of 741 nm selected by an interference filter. The setup allowed lateral resolution of the surface enabling simultaneous monitoring of interactions on different regions of the chip.

The ellipsometric mode, where both the intensity and the phase changes of the reflected light are monitored and converted into two ellipsometric angles Ψ and Δ, respectively. For the parameter set used in our experiments, Δ is inversely proportional to the mass adsorbed to the gold chip, while Ψ is directly proportional to it. The angle of incidence was 70° in ellipsometric mode and the so-called “fixed compensator nulling scheme” was used, where the compensator angle is set at 45° and the polarizer and analyzer are rotated for the measurement. The regions of interest were selected for the relevant patches of the array, reference regions of interest were selected for non-patterned background, and the averaged signals in each region were monitored over time. For the kinetic mode the sampling rate was 10 s.

### SPR measurements

The SPR surfaces consisted of 12 mm×12 mm glass slides coated with a 45nm thick film of gold. Prior to use, SPR surfaces were modified with surface active molecule ODT (octadecanethiol; C_18_ ; HSCH_17_-CH_3_). ODT was transferred to the gold surface using microcontact printing polydimethylsiloxane (PDMS) stamps with line and square features. Conditions for PDMS stamp preparation and for microcontact printing are described elsewhere.

For kinetic experiments the following protocol has been followed: as illustrated in fig. 2, after multistep preparation of the gold chip (see below), TeNT-46 was injected into the SPR cell under continuous flow (at a rate of 30 μl/min), and the signal was equilibrated with HBSS/10 mM HEPES running buffer, pH 6.5. After saturation of the SPR signal, buffer was allowed to flow in to rinse off unbound probe.

#### Gold chip preparation

Prior to tethering of the ODT layer, the equipment and the chips were cleaned in RCA solution (5:1:1 mixture of MilliQ de-ionized (MQ) water, hydrogen peroxide and 25% ammonium) for 5 minutes at 80°C and afterwards sonicated in MQ water for another 5 minutes. The PDMS stamps were sonicated in absolute ethanol for 3 minutes, then dried under a nitrogen stream and incubated with 100 µL of ODT solution for 5 minutes. Afterwards, the incubation the stamp was blow dried and gently pressed onto a cleaned and dried gold chip for 10 minutes. The gold chip was then sonicated in absolute ethanol for 1 minute. The SPR chamber and tubing were cleaned by rinsing with RCA solution at 80 °C and with MilliQ distilled water. A completely blow dried chip was gently placed onto the flow chamber with the tethered gold side facing down. A drop of refractive index matching oil (N_D_ = 1.5167 ± 0.0005; Cargille Labs) was then applied on the chip before the glass prism was mounted.

### Vesicle formation

Vesicles were made from a core ternary lipid mixture of POPC:SM:Chol (1:1:1) with an addition of GT1b at varying concentration of 0%, 1%, 5%, 10%, 15% and 20% (mol-%). The required composition was calculated and mixed in a round bottom flask to a final mass of 1 mg. Using an evaporator, the solvents were removed under a vacuum and at 65 °C for 2 hours to allow the lipids to dry. Dry lipids were resuspended in buffer (HBSS/10mM HEPES buffer at pH 6.5). The mixture was thereafter extruded to form LUVs as described in (Mayer et al., 1986). Briefly, the mixture was frozen and then thawed above transition temperature (65 °C) in 5 cycles and then passed through a 50 nm polycarbonate membrane (Whatman) 23 times, using an Avestin Lipofast extruder, resulting in vesicles with a diameter of about 80 nm. Vesicles were used within 24 hours.

For the surface characterization with polarization modulation-infrared reflection absorption spectroscopy (PM-IRRAS), a silicon wafer with an evaporated 20 nm gold coating was cleaned in a RCA bath for 8 minutes at 80°C. The slide was then rinsed and sonicated in MQ-water for another 5 minutes. A 10 mM ODT-solution was incubated on the surface in a dark cabinet overnight. Afterwards, the slides were sonicated for another 3 minutes in pure ethanol and dried under a stream of nitrogen. In order to protonate the slides for the PM-IRRAS measurement, they were placed into a solution of 0.5M NaCl in water for 30 minutes and then dried again with a nitrogen stream. To monitor the formation of additional peaks during the spread of vesicles onto the ODT surface, the previously measured chips were incubated with a solution of POPC/SM/C (1:1:1) and 5% GT1b in HBSS/10 mM HEPES buffer pH 6.5 for one hour, rinsed with HBSS/10 mM HEPES buffer pH 7.5 for 5 minutes, and dried under a nitrogen stream.

Giant unilamellar vesicles (GUVs) were formed by mixing the lipid components with 0.1% Rhodamine-DPPE (lissamine rhodamine B 1,2-dihexadecanoyl-*sn*-glycero-3-phosphoethanolamine, triethylammonium salt; Rho-DHPE) and imaged with a Deltavision wide-field microscope (Applied Precision). The components were dried, and a 200 mM sucrose solution was added. GUVs were produced by electroformation with a Nanion prep pro (Nanion Technologies GmbH) at 5 Hz, and 13 V, at 45°C for 2 hours. Other details of GUV formation and imaging are described in Oglęcka et al (Oglecka et al., 2014).

### Hybrid lipid bilayer formation

The ODT-stamped gold chip was mounted in the iSPR chamber and rinsed with HBSS/10mM HEPES buffer at pH 6.5. Thereafter, large unilamellar vesicles (LUVs) of the desired composition were applied and fused with calcium-containing buffer. After 1 hour of incubation the chamber was rinsed with HBSS/10mM HEPES buffer at pH 7.5, causing non-fused vesicles to rupture.

Images of the hybrid bilayer on the gold surface were taken with a DeltaVision (Applied Precision) wide-field microscope using a 10x Olympus lens and air-cooled CCD camera. Images were collected with SoftWorx software (Applied Precision), images and processed with ImageJ.

### Polarized modulation-infrared reflection-adsorption spectroscopy (PM-IRRAS)

Film properties of the membrane attached to the gold substrate were characterized with Polarization Modulation Accessory 50 (PMA 50) from Bruker (Billerica, MA). The detector angle was set to 83° and the sample was purged with a stream of nitrogen (flow rate: 10 L/min). Data were recorded from 850 to 4000 cm^-1^, with a resolution of 4 cm^-1^ and a sample scan time of 5 mins. After data collection, the baseline for each spectrum was calculated by a polynomial fitting function.

### Fluorescence Correlation Spectroscopy (FCS)

FCS experiments were done on an Olympus FluoView 300 confocal microscope with the custom-built FCS detection unit on top of the scanning unit. A 543 nm helium-neon laser was used to excite the TAMRA fluorophores and the laser power was maintained at 25 µW for all experiments. Details of the instrumentation are described in (Hebbar et al., 2008). The fluorescence signals were collected onto a single photon sensitive avalanche photo diode (APD; SPCM-AQR-14, Pacer Components, UK) after passing through an emission filter 595 AF60 (Omega Optical Inc., Brattleboro, VT) in point-scanning mode, and fitting of autocorrelation curves was done as described (Hebbar et al., 2008) using a self-written program in IgorPro 6.0 (Wavemetrics, Portland, OR) with 2D or 3D and 1- or 2-particle, 1 triplet models, yielding diffusion times τD1 and τD2 and corresponding mole-fractions F1 and F2. FCS correlation curves were fitted to 3D, one particle (corresponding to freely diffusing, unbound probe), and 2D, 2 particle (corresponding to membrane-bound, bimodally diffusing) models, as follows:

3D1P1t:

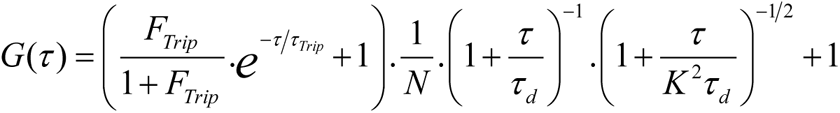

2D2P1t:

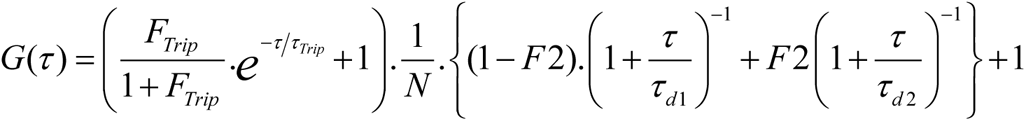

Here *N* refers to the number of particles in the observation volume; *τ_D1_* and *τ_D2_* are the characteristic diffusion times of different particles; the mole fraction of particle 2 is given by *F_2_= 1-F_1_*; the fraction of molecules in the triplet state and the characteristic triplet state time are given by *F_trip_* and *τ_trip_*, respectively.

## Results

### Design of a ganglioside binding tetanus-derived fluorescent peptide probe, TeNT46

In order to examine the nature of membrane glycolipid-containing, toxin-interacting domains in living cell membranes, we designed a short (46aa) peptide probe derived from the heavy chain (Hc) domain of tetanus, a clostridial toxin known to bind gangliosides abundant on neuronal membranes, including GT1b (Halpern and Loftus, 1993). We chose tetanus toxin as a starting molecule because its binding to gangliosides has been characterized in detail at the molecular and structural level, with the C-terminal end of the tetanus Hc mediating the interaction with gangliosides (Chen et al., 2009; Emsley et al., 2000; Fotinou et al., 2001; Ginalski et al., 2000; Rummel et al., 2003). The defined lipid-interacting region at the C-terminal tail of the Hc domain provided us with a reasonable guide toward the design of a small glycolipid-targeted membrane domain-binding probe owing to its well-characterized interaction with gangliosides, including GT1b, GD1b, GM1a, and GD3.

The C-terminal-most residues of the tetanus Hc β-trefoil domain, (amino acids 1110-1315, termed H_CC_), contains two distinct regions determined by both mutation analysis and X-ray crystallographic structure to bind to, respectively, the headgroup core region (the so-called R pocket, or lactose binding site) and the sialic acid residues of GT1b (the W pocket) (Chen et al., 2009; Emsley et al., 2000; Fotinou et al., 2001; Rummel et al., 2003). TeNT46 peptide, encompassing residues 1270-1315 of H_CC_, includes six of the nine amino acids within the C-terminally located lactic acid binding domain whose mutation impairs binding (the region is shown boxed in fig. 1A (Rummel et al., 2003); the 46 aa C-terminal-most segment of it that constitutes TeNT46 is modelled in fig. 1B). This 46aa peptide was linked to a TAMRA fluorophore at the N-terminus via two copies of an amino-polyethyleneglycol (PEG) linker (fig. 1C; described in (Steinert et al., 2008)).

**Figure 1.**
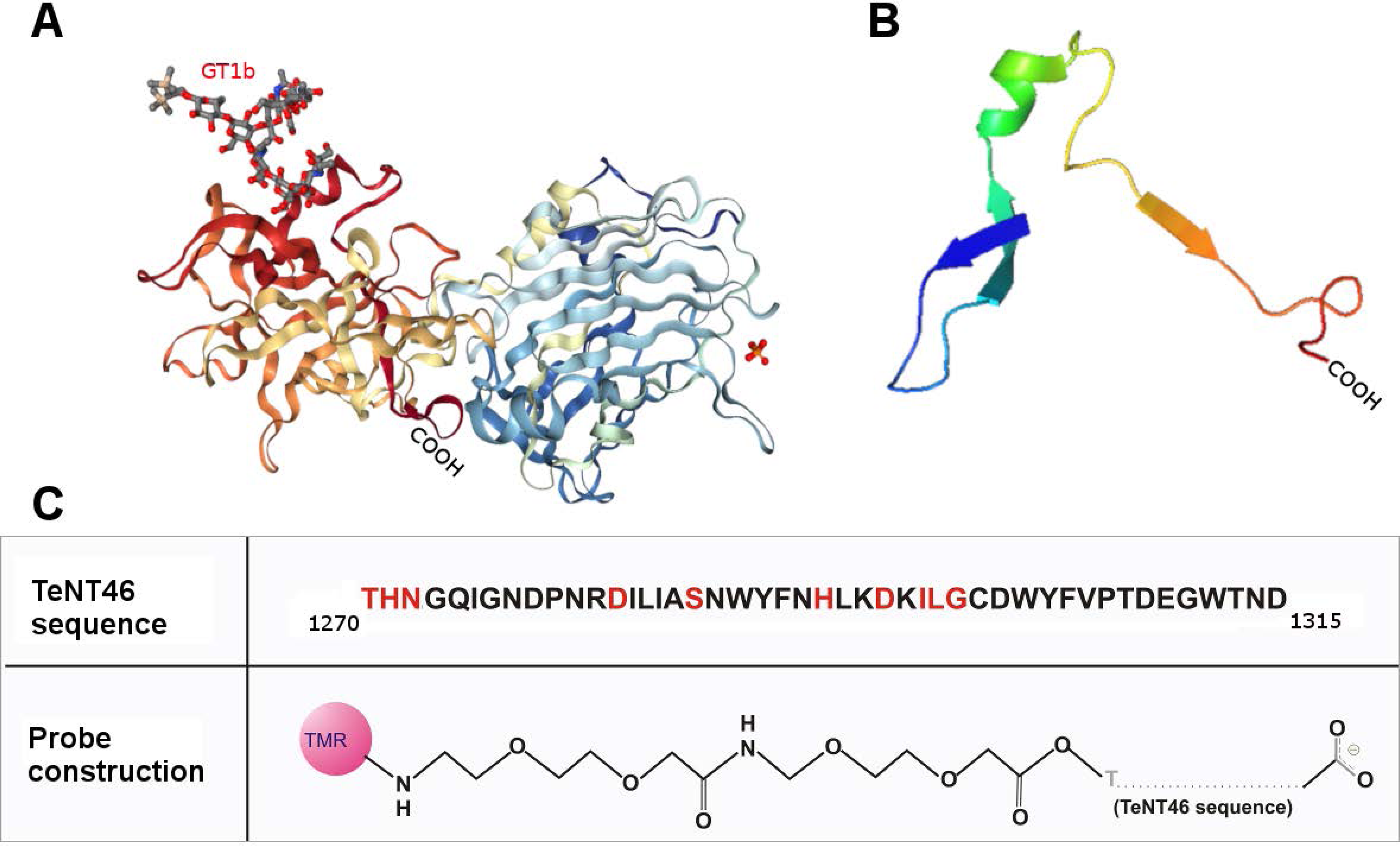
Design of the tetanus-derived probe, TeNT46. A) Crystallographic structure of the tetanus toxin heavy chain (Hc) fragment as a ribbon model, showing the GT1b (stick model) binding site (Protein Databank No. 1FV3 (Berman et al., 2000)(Fotinou et al., 2001)), most of which is encompassed in the TeNT46 sequence; B) ribbon model of the TeNT46 peptide alone, generated by truncating the above crystallographic structure of tetanus Hc at residue 1270; C) Amino acid sequence of TeNT46 peptide, showing conserved residues that participate in ganglioside binding, and probe construction, consisting of a tetramethylrhodamine (TMR) fluorophore linked via a double amino-PEG (AEEAc) linker at the amino end to the C-terminal-most 46 amino acids of the Hc domain.

**Figure 2.**
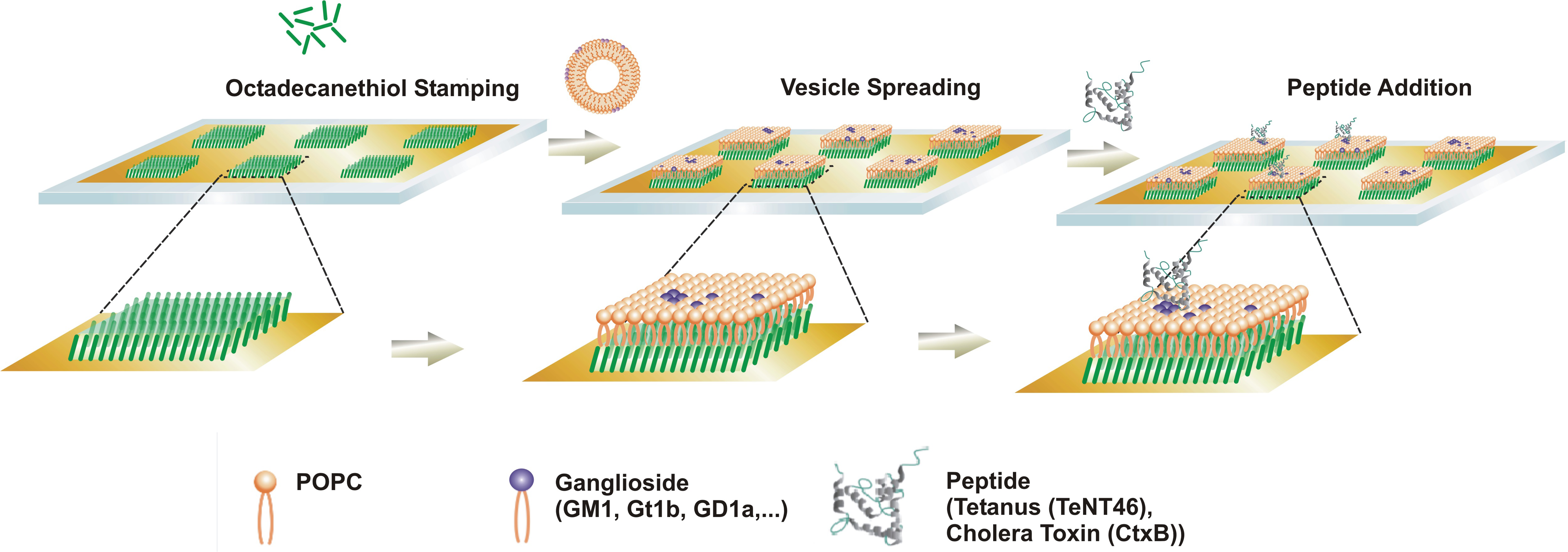
Fabrication steps of C18 ODT tethered lipid monolayer platform and illustration of proposed methodology for TeNT-46 detection.

This linker construction was earlier shown to interfere minimally with binding of another ganglioside-interacting probe, the Sphingolipid Binding Domain of Aβ (SBD; (Hebbar et al., 2008; Lauterbach et al., 2012; Steinert et al., 2008; Wang et al., 2015)). The addition of the fluorophore allowed us to characterize the diffusion behaviour of the peptide via spectroscopic and fluorescence imaging, in parallel with the imaging-surface plasmon resonance (iSPR) binding studies described in the next section.

### A hybrid lipid bilayer as a biosensing surface for peptide probes

To determine whether the TeNT46 peptide probe was able to bind to lipid surfaces, and to find out whether the expected ganglioside binding preferences of the parent toxin would be preserved, we first used iSPR with a variety of plasma membrane-like lipid substrates for the hybrid supported lipid bilayer (SLB) to assay binding and lipid specificity.

Commonly encountered problems with standard SLB deposition onto the sensor chip used in SPR are weak attachment and non-fusion of the applied liposomes. Especially problematic are liposomes with a high proportion of charged lipids, e.g. gangliosides, such as the mixtures used in this study. To overcome these problems, a hybrid bilayer was constructed in a 2-step process, wherein the bottom layer (in contact with oxidized gold) was formed as a flat layer, stabilized by covalent thioester attachment of the lipid acyl chain-like octadecanethiol (ODT), and patterned onto the surface by means of a PDMS stamper (fig. 2). This formed the “inner leaflet” of the resulting hybrid bilayer. The upper layer (in contact with the analyte, analogous to the outer leaflet of the plasma membrane) was generated from 100 nm liposomes (large unilamellar vesicles; LUVs) consisting of a plasma membrane-like mixture of POPC/SM/Chol (1:1:1) and 5% ganglioside, deposited onto the ODT layer and fused by raising the pH of the buffer from 6.5 to 7.5 (see Methods).

iSPR provided visual verification of the substrate surface as it was laid down layer-by-layer. After deposition of a patterned lipid substrate of the desired composition onto a gold chip, the thickness of the substrate was determined ellipsometrically, and binding of an analyte spread over this surface was then measured by SPR.

The angstrom-resolution surface height-imaging capability of the iSPR setup was crucial for the ability to detect formation of the hybrid bilayer surface, which is expected to be on the order of 4-5 nm. A further important feature of the iSPR platform used here was the patterning of the hybrid bilayer surface. Below, we describe a novel variation of microcontact printing, where a hybrid lipid bilayer substrate is patterned onto a gold chip.

The surface was monitored at each step of the deposition by ellipsometric imaging via the CCD camera in the iSPR setup (Accurion GmbH). To do this, several regions of interest (ROIs) were defined, first comparing ODT-stamped patches with bare gold inter-patch areas to determine the height of the presumptive C_18_ layer formed from the ODT, before vesicle addition. Keeping the regions of interest unchanged, LUVs were eluted to complete the formation of a hybrid bilayer on patches, and the heights of the previously selected points were measured during deposition. The ODT attachment step yielded a height increase of ∼30Å (see fig. 3A), while LUV addition caused a sudden increase, with a subsequent drop in signal (Δ). These height changes could be explained by the initial docking of vesicles on C_18_ patches, leading to a substantial height increase, followed by rupture and spreading to form the upper half of a hybrid bilayer. Thickness measurement of this upper layer yielded a 21 Å increase in height (fig. 3B). The detected height increase of 2 – 2.5 nm for a monolayer is consistent with an overall membrane bilayer thickness of 4 – 5 nm, corresponding to the expected height of a fused hybrid bilayer.

**Figure 3.**
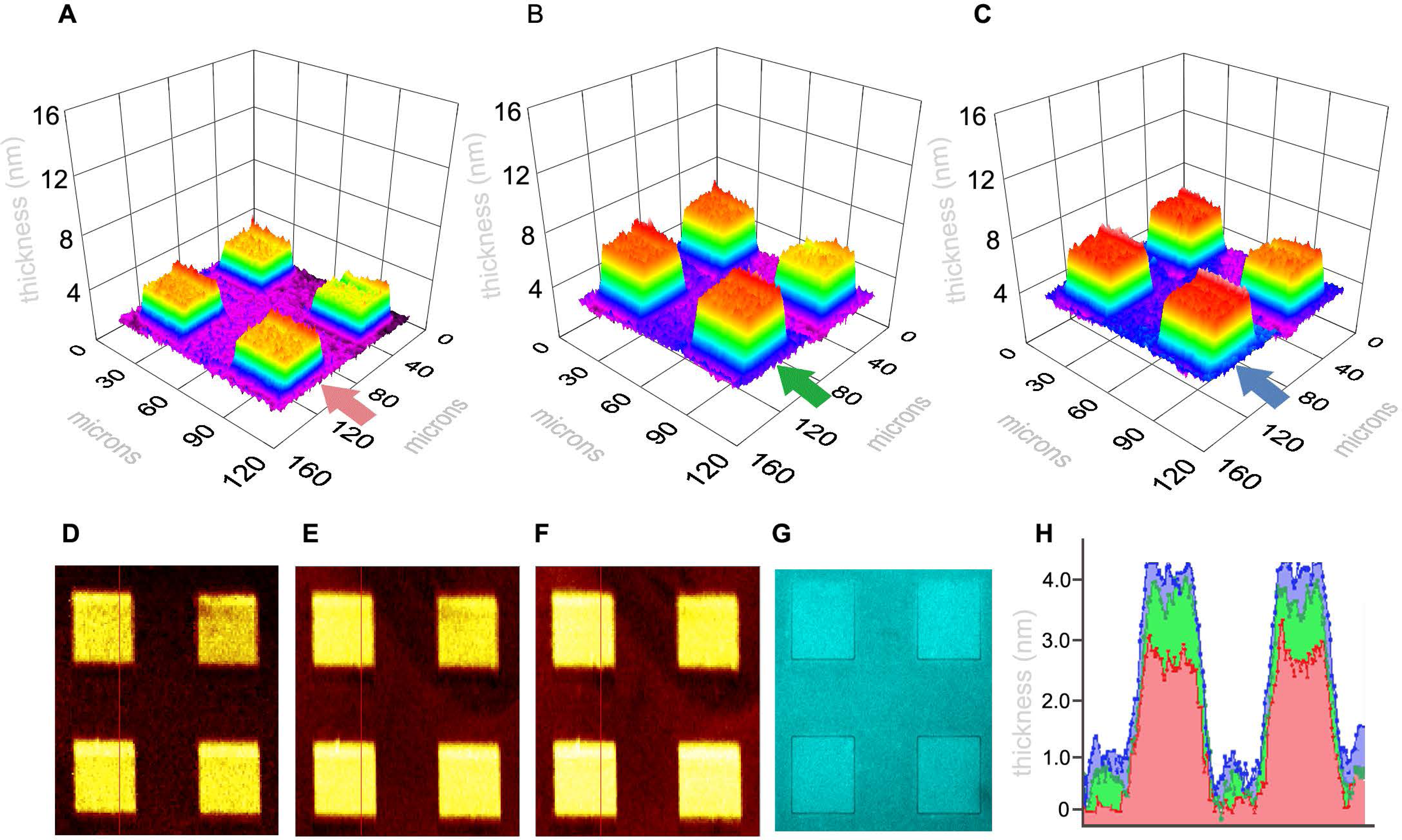
iSPR height maps during layer-by-layer construction of the OTD-lipid surface are consistent with formation of a bilayer. A)-C) 3-dimensional maps and height profile of C18 modified chip, A) before LUV addition, B) after LUV addition and rupture, C) after TeNT46 incubation. D)-G) show the respective real-time images of the surfaces. H) height map before (red) and after (green) addition and rupture of LUVs and (blue) the injection and incubation of TeNT46.

After the addition and incubation of the TeNT46 derived toxin, only a small increase in height was detected (fig. 3C). This is expected, given the concentration of the peptide and the likelihood of a low degree of coverage of peptide on the bilayer surface.

Fabrication of the 21Å smooth C_18_ supported monolayer array was consistently reproducible, and the 3D map also revealed a “smooth topography”, implying full or nearly full coverage of lipid monolayer, as far as the limits of the lateral resolution of the iSPR instrument allow. The gold surface was imaged after application of the hybrid bilayer by widefield microscopy. The fluorescence image (fig. 4A) shows that the stamped ODT patches (squares), which should only carry labelled DHPE in the upper leaflet of the hybrid bilayer, appear darker than the surrounding non-stamped area. We assume that this is due to the attachment of intact vesicles to the bare gold substrate between the patches, as opposed to the ruptured monolayer on top of the ODT-patches, resulting in a brighter fluorescent signal.

**Figure 4.**
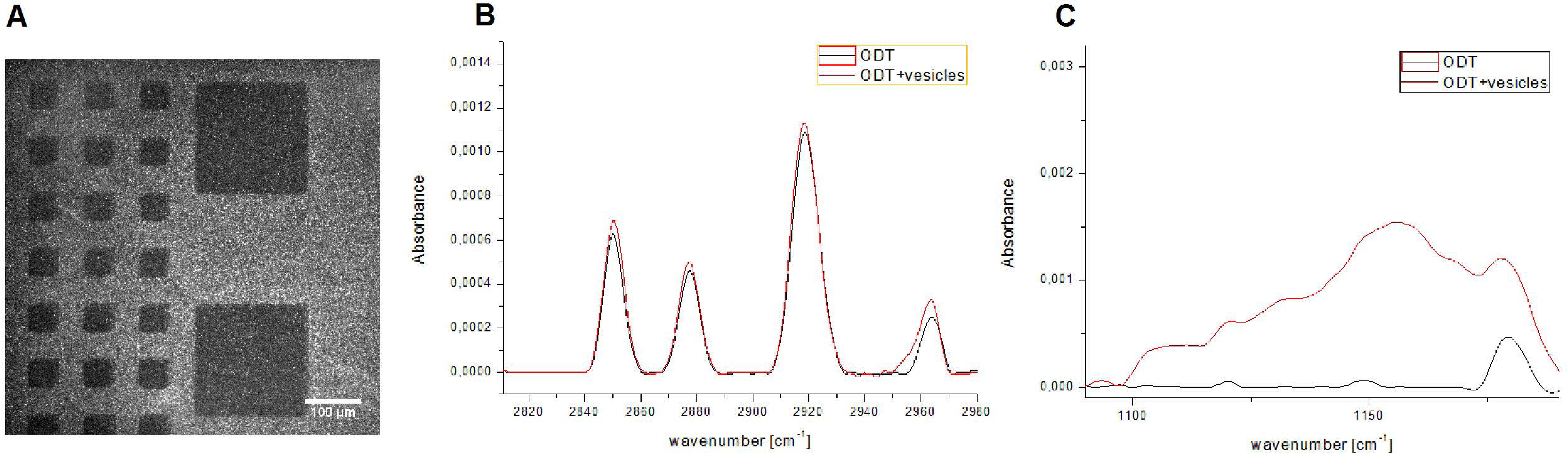
A) Fluorescence image of the stamped hybrid bilayer surface on the gold chip. Stamped areas appear darker; B) PM-IRRAS monitoring of the formation of the hybrid bilayer. The region 2820-2980 cm^-1^ shows CH_2_ and CH_3_ modes of the ODT monolayer and the lipid chains in the other leaflet of the hybrid bilayer. C) Fingerprint region before and after spreading of lipid vesicles composed of POPC/SM/C (1:1:1) and 5 % GT1b on the ODT SAM.

### Characterization of the formation of a self-assembled monolayer

During the stamping process, a self-assembled monolayer of ODT molecules is transferred from the PDMS stamp to the gold surface (Plant, 1999). In order to obtain reproducible binding experiments, the self-assembled ODT monolayer should be of high quality in terms of molecular packing and in-plane organization/uniformity. Herein we used PM-IRRAS to characterize the ODT monolayer in the CH_2_ and CH_3_ stretching and fingerprint regions of the infrared spectrum. Two peaks are clearly visible at 2850 cm^-1^ and 2918 cm^-1^, and they are characteristic for symmetric and asymmetric CH_2_ modes in a highly ordered and densely packed *all trans* assembly of alkyl chains (Ramin et al., 2012). Additionally, peaks at 2877 cm^-1^ and 2964 cm^-1^ are also seen and they correspond to the symmetric and asymmetric stretching modes of the terminal CH_3_ group (Dilimon et al., 2013). After adding a solution of vesicles and proceeding with hybrid lipid bilayer formation as described in fig. 3, the sample is characterized again with PM-IRRAS. Comparing the spectra before and after vesicle spreading (black and red lines in fig. 4B and C), the peaks for the CH_2_ stretching modes are slightly shifted to higher wavenumbers (e.g. from 2918 cm^-1^ to 2920 cm^-1^for the asymmetric CH_2_ mode) and the corresponding intensities are increased. We attribute this to the deposition of the upper half of the hybrid lipid monolayer on top of the ODT SAM, increasing the number of CH_2_ groups and therefore the area under the peaks. The slight shift in the CH_2_ modes can be explained by a more loosely formed, less organized second lipid layer.

Fig. 4C shows the fingerprint region of the spectra of the ODT monolayer and the hybrid bilayer. The increase in peak intensities in this region (red curve) derives from the terminal part of the lipids, in this case POPC/SM/C (1:1:1) and 5 % GT1b. All the lipids in this mixture contain chemical groups with characteristic vibrational frequencies in this region (Brandenburg and Seydel, 2014; Krafft et al., 2005). However, individual peaks in the spectra cannot provide a “fingerprint” for single lipid species, due to the presence of different lipids with a variety of chemical groups and the complex shape of the IR spectrum. For instance, ester groups, palmitic acid and phospholipid bands from POPC (1122, 1126, 1129, 1174 cm^-1^) and acyl chains from sphingomyelin all appear in this region (1128 cm^-1^). Furthermore, owing to its complex structure, the ganglioside GT1b contains a lipid component, a ceramide backbone and sugar residues, that also absorb strongly in this range. Specifically, the acyl chain absorbs at 1130 cm^-1^ and the sugar rings inter alia at 1117 cm^-1^. We conclude from the PM-IRRAS experiments that a uniform and densely packed ODT monolayer is formed, with a less organized POPC/SM/Chol/GT1b mixed monolayer on top of this ODT SAM.

### iSPR with ODT-ganglioside hybrid bilayers confirm binding of CTxB to GM1

Before exploring possible binding targets of the novel probe, TeNT46, we first wished to test the ability of the iSPR platform with a hybrid ODT-lipid bilayer to detect binding of the well-studied microdomain and ganglioside GM1-binding probe, the B subunit of Cholera toxin (CTxB) (Holmgren et al., 1975; Milani et al., 1992) to its preferred lipid target.

The target membrane (i.e. the top leaflet of the hybrid bilayer) consisted of POPC/SM/Chol (1:1:1) liposomes with 5% of one of five gangliosides (GT1b, GD1a, GQ1b, GD1b, or GM1) (fig. 5A), or varying amounts of GM1 ganglioside (fig. 5B). The usual blocking step of the surface (Vashist, 2011; Vollmer et al., 2015) with BSA was eliminated, since measurements of binding were made from sites within the ODT patches, with reference to binding in inter-patch regions (see fig. 3D-G). By avoiding the BSA blocking step, possible blocking agent artefacts on the hydrophobic patches have also been eliminated.

**Figure 5.**
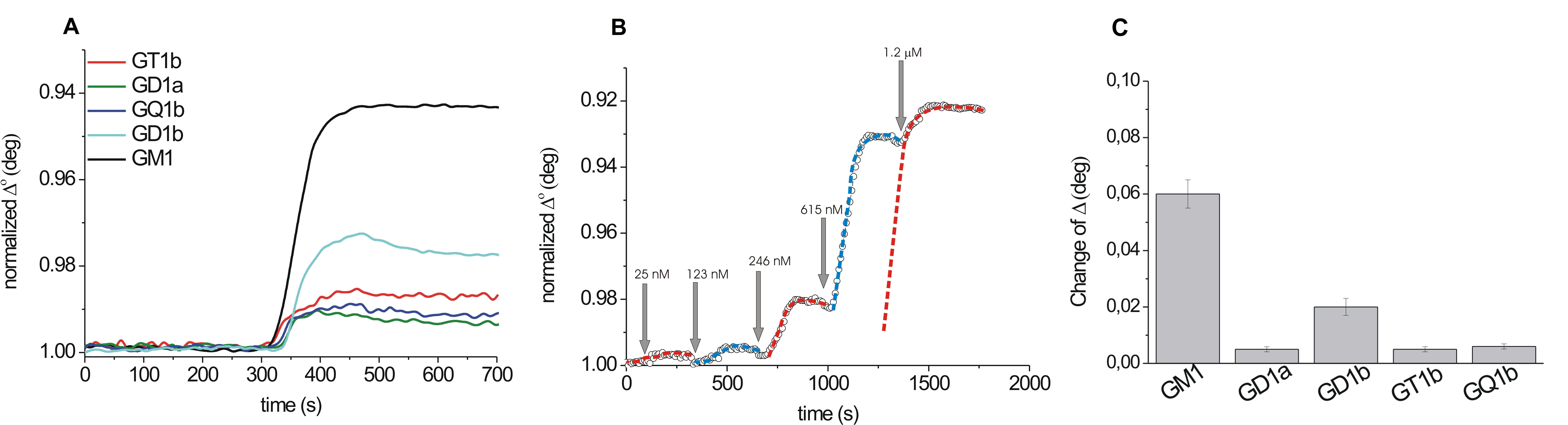
iSPR measurements showing change in the ellipsometric angle (indicative of binding) during flow-through of 1 uM CTxB onto hybrid bilayers. An upper layer consisting of 5% of different gangliosides in a 1:1:1 mixture of POPC/Chol/SM detects (A) specificity for GM1 over other gangliosides; (B) binding curves at different concentrations of CTxB. (C) Average magnitude of the change in ellipsometric angle induced by CTxB on bilayers containing different gangliosides.

Of the gangliosides tested, including GT1b, GQ1b, GD1b and GD1a, and GM1, CTxB bound preferentially to GM1 (fig. 5) matching the known lipid specificity for this toxin. Importantly, the stamping of a patterned bilayer enabled many simultaneous measurements to be carried out at multiple points on a single surface, with the inter-patch regions presenting an intrinsic internal reference. This strategy greatly improved the statistical reliability of the method, since >40 measurements could be averaged for each composition. Recovery of GM1 as a receptor for CTxB, with much stronger relative affinity than other gangliosides tested, agreed with published data (Kuziemko et al., 1996; MacKenzie et al., 1997; Seo et al., 2013), and demonstrated the validity of the platform for measuring peptide-lipid interactions.

### TeNT46 is a novel GT1b-binding tetanus toxin-derived probe

Having validated the effectiveness of the iSPR/hybrid bilayer platform with CTxB, we wanted to test TeNT46 binding to various ganglioside-containing surfaces in the same manner. TeNT46 peptide was expected to differ in ganglioside specificity from CTxB, as a substantial literature has established GT1b as the major target for attachment of the complete toxin to neuronal membranes (Emsley et al., 2000; Fotinou et al., 2001; Louch et al., 2002; Rummel et al., 2003), although other gangliosides such as GD1a, GD1b, and GD3 have also been identified as potential receptors (Chen et al., 2009; Deinhardt et al., 2006; MacKenzie et al., 1997; Masuyer et al., 2017; Singh et al., 2000).

To monitor TeNT46 peptide binding, we applied 5% ganglioside-containing liposomes to create the bilayer upper leaflet, similarly to the experiments with CTxB. Testing of GM1, GD1a, GD1b, GQ1b, and GT1b confirmed that among those, GT1b was indeed the preferred binding target of this TeNT46 fragment, in agreement with the known preference of the intact tetanus toxin (fig. 6). Varying the GT1b content of the upper leaflet from 0 to 20 mol% showed a steady increase in binding of TeNT46 to the surface (fig. 6C).

**Figure 6.**
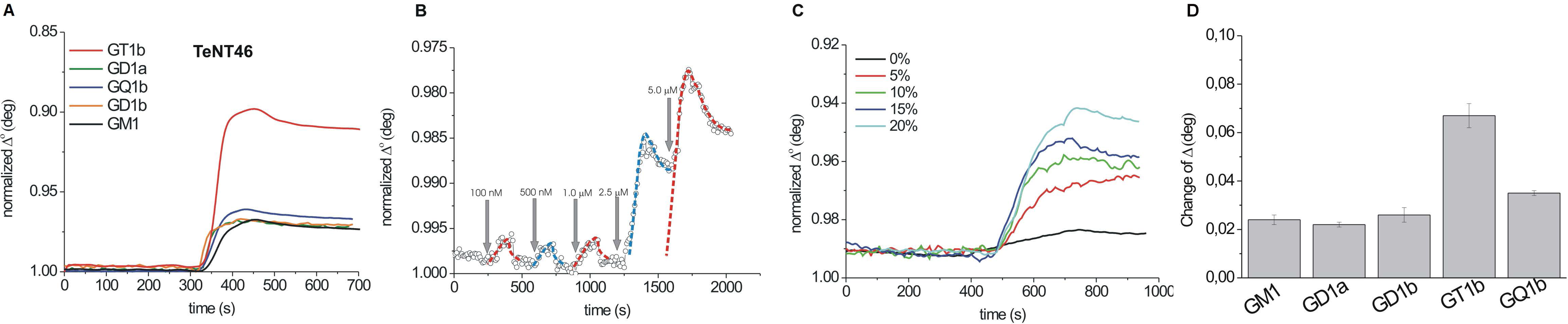
iSPR reveals specificity of TeNT46 probe for ganglioside GT1b. A) iSPR measurements showing change in the ellipsometric angle (indicative of binding) during flow-through of 5 uM TeNT46 onto hybrid bilayers. An upper layer consisting of 5% of different ganglioside in a 1:1:1 mixture of POPC/Chol/SM detects (A) specificity for GT1b over other gangliosides; B) binding curves at different concentrations of TeNT46; C) Average magnitude of the change in ellipsometric angle induced by TeNT46 on bilayers containing different gangliosides..

### Different lipid binding requirements of CTxB and TeNT46 in the iSPR platform predict differences in diffusion behaviour on cells

In order to test the non-ganglioside lipid requirements of CTxB and TeNT46 binding, cholesterol, SM, and ganglioside were each removed singly or in combination from the POPC/SM/Chol/ganglioside lipid mixtures used to make the upper surface of the hybrid bilayer. For CTxB, as expected (Hebbar et al., 2008; Lingwood et al., 2008), cholesterol was required for optimum binding, whereas SM was not required, and was even slightly inhibitory (fig. 7A, B). Conversely, TeNT46 did not require cholesterol, which appeared to have an inhibitory effect on binding, particularly in the absence of SM. Unlike CTxB, it absolutely required SM (fig. 7C, D).

**Figure 7.**
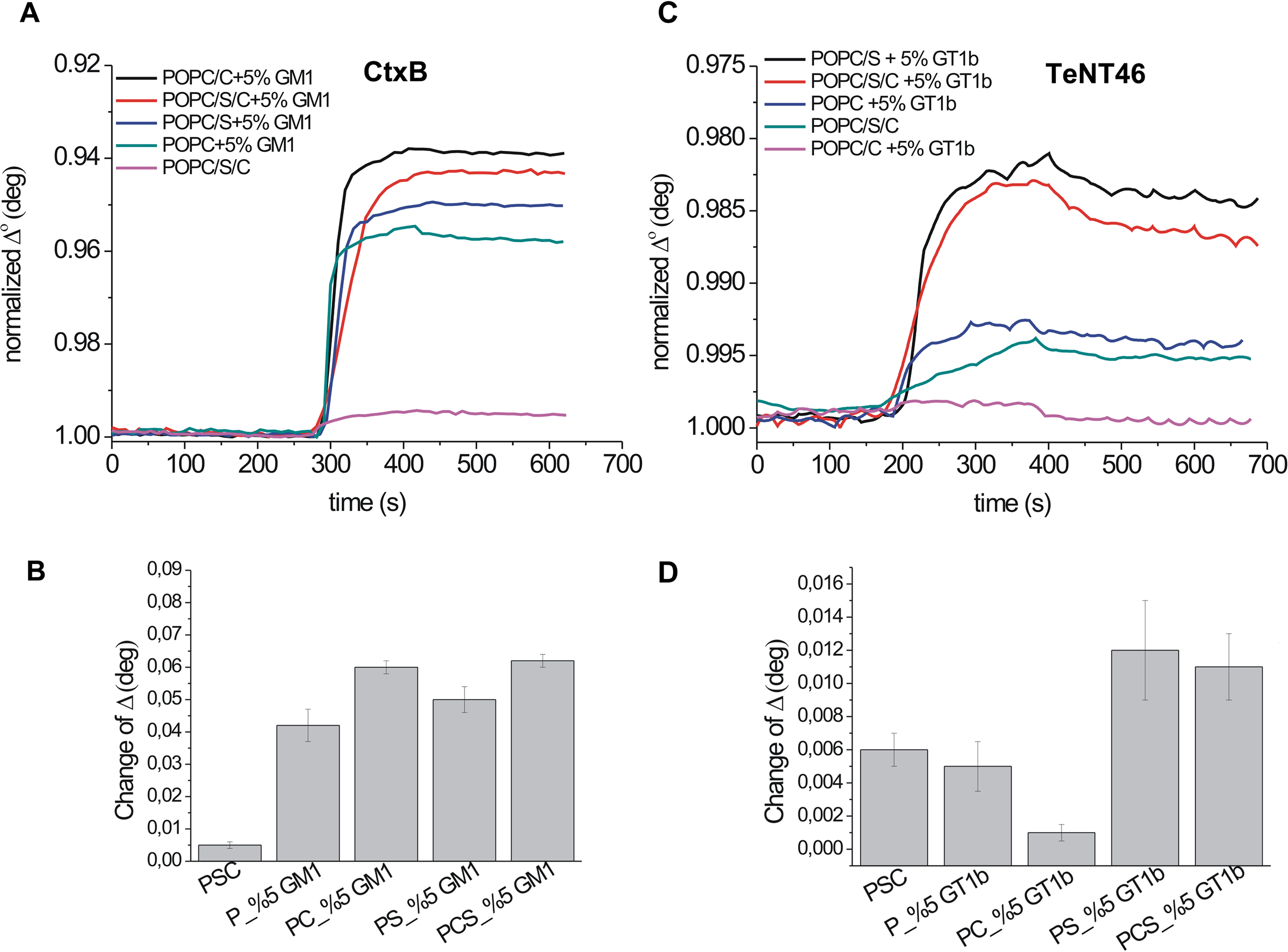
CTxB and TeNT46 prefer different constellations of lipids, as shown by differing sensitivities to the removal of cholesterol and sphingomyelin. A) iSPR measurements show CTxB binding to membranes containing GM1 and either a complete complement of a plasma membrane-like mixture (POPC/Chol/SM) or with individual components missing; B) quantification of the change in ellipsometric angle (indicative of binding) during flow-through of CTxB on the different surfaces; C) TeNT46 binding to similar combinations of lipids with its target ganglioside, GT1b; D) quantification of TeNT46 binding, as in B).

In an effort to determine whether our findings in the iSPR system corresponded to lipid preferences and binding behaviour on actual cell membranes, we examined the diffusion characteristics of the two peptides, CTxB and TeNT46 on the plasma membrane of live neuroblastoma SH-SY5Y cells by confocal fluorescence correlation spectroscopy (FCS). All data were optimally fit with a two-particle model in which the first component had a diffusion time τD1 < 1 ms, while the second particle had an average diffusion time of τD2 > 10 ms. The distribution of diffusion times recovered for τD2 are shown as histograms for each of the peptides (figs. 8A, 9A). The τD1 values (not shown in the histogram) are presumably from probe measured in solution, since these values are about an order of magnitude faster than the τD2 values, which we took as the diffusion times of the membrane bound peptides, and are typical for membrane-bound probe (Garcia-Saez and Schwille, 2010).

**Figure 8.**
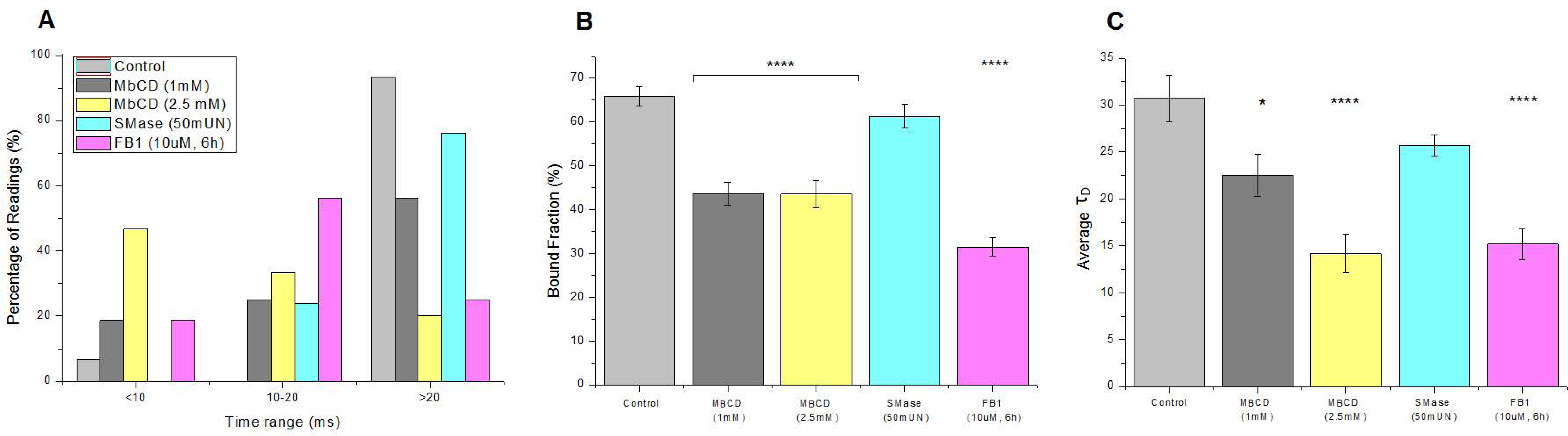
CTxB diffusion and binding assayed by FCS show sensitivity to cholesterol and sphingolipid depletion, but not to sphingomyelinase (SMase). A) Diffusion times (τD) of CTxB (10ug/ml) on plasma membrane of SH-SY5Y in control vehicle-treated cells, and after cholesterol extraction with mβCD, bacterial SMase, or ceramide synthase inhibitor FB1 treatment, shown as histograms that shift leftward as diffusion rate increases; (B) bound fraction (percentage of reads fitting F2) shows reduction of membrane-bound CTxB after mβCD and FB1, but not SMase; C) Average diffusion times shift significantly after removal of cholesterol and sphingolipids with FB1. Bars indicate SEM from at least 15 independent measurements, each of which is the result of 1 minute of continuous data collection.

**Figure 9.**
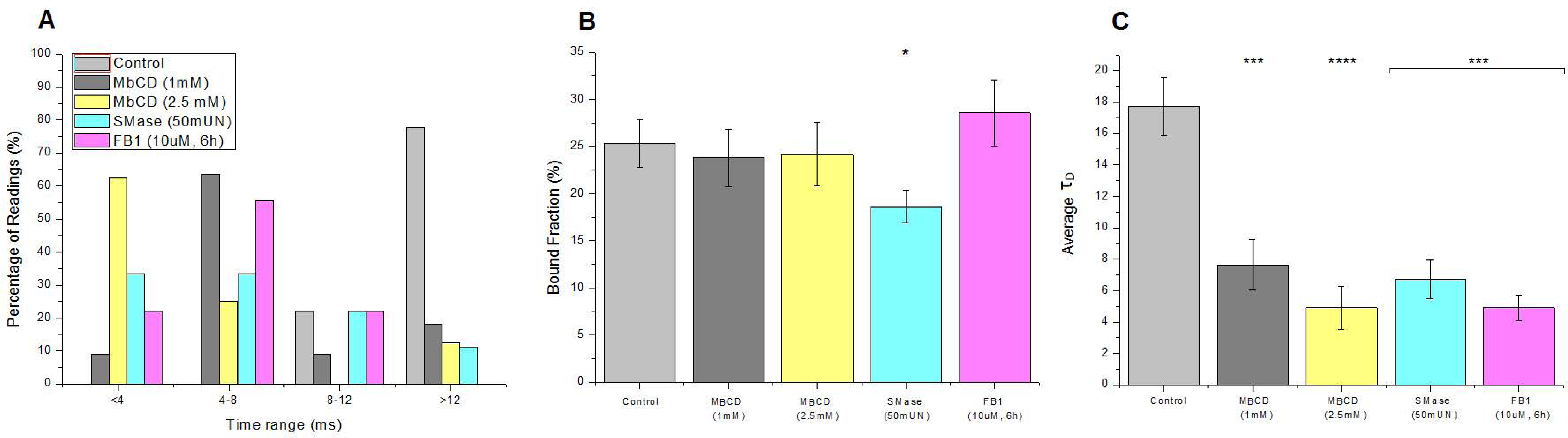
TeNT46 diffusion rate is highly sensitive to cholesterol and sphingolipid depletion, whereas binding is only diminished significantly by sphingomyelin loss. A) Diffusion times (τD) of TeNT46 (10ug/ml) on plasma membrane of SH-SY5Y in control vehicle-treated cells, and after cholesterol extraction with mβCD, bacterial sphingomyelinase (SMase), or ceramide synthase inhibitor FB1 treatment, shown as histograms that shift leftward as diffusion rate increases. B) Bound fraction (percentage of reads fitting F2) shows a significant reduction of membrane-bound TeNT46 only after SMase treatment. C) Average diffusion times shorten significantly after removal of cholesterol, sphingomyelin, or sphingolipids with FB1. Bars indicate SEM from at least 15 independent measurements, each of which is the result of 1 minute of continuous data collection.

Strikingly, changes in diffusion behaviour of CTxB and TeNT46 after lipid depletions of cells mirrored their differing requirements for cholesterol vs. SM in the iSPR experiments presented above. On neuroblastoma SH-SY5Y, diffusion differed between CTxB and TeNT46, with average diffusion times τD2 of 30.7 ± 2.5 ms and 17.7 ± 1.8 ms respectively (figs 8, 9). These diffusion times are both faster than the diffusion we reported earlier for another ganglioside-interacting probe, SBD, which had a τD2 of ∼60 ms on the same cell type (Hebbar et al., 2008; Sankaran et al., 2009).

In parallel with the iSPR binding studies, we also wanted to use FCS to test which canonical lipid components of ordered domains--cholesterol, sphingomyelin, or sphingolipids in general--influenced the membrane association and the diffusion behaviours of the two probes. Our expectation based on published work (Bacia et al., 2004; Hebbar et al., 2008; Lingwood et al., 2008; Šachl et al., 2015; van Zanten et al., 2010) was that cholesterol and sphingolipid depletion should affect CTxB diffusion and binding, but that sphingomyelin may not (Stefl et al., 2012). Cholesterol depletion with 1 mM mβCD--a comparatively low dose (Chadda et al., 2007; Saha et al., 2015), but sufficient in our experience to abolish the slowly diffusing component of another ordered domain probe (Kraut et al., 2012)(M. Manna PhD thesis, National Univ. Singapore)-- indeed significantly decreased the fraction of CTxB bound from 65.8% to ∼43.5%, the same decrease as treatment with 2.5 mM mβCD. These concentrations of mβCD progressively shortened the slow τD of the probe from 30.7 ms to 22.52 ms and 14.19 ms, respectively (fig. 8C). The sphingomyelin synthase inhibitor Fumonisin B1 (FB1), which suppresses synthesis of sphingolipids by inhibiting sphingosine N-acyl transferase (Merrill et al., 1993), also reduced the fraction bound to 31.5% (fig. 8B). SMase, however, had no significant effect on average τD or fraction bound (25.69 ms and 61.4%), although the histogram appeared to shift slightly toward shorter transit times (fig. 8A, C). The conclusion we draw from these results is that CTxB probe, as expected, requires cholesterol, but not SM, for both optimum binding and slow diffusion.

Strikingly different results were seen for TeNT46 under mβCD cholesterol extraction: the slowly-diffusing fraction was, similarly to CTxB, strongly released from confined diffusion (fig. 9A, C), indicated by the leftward shift in the τD histogram and shortening of τD from 17.72 ms to 7.64 ms and 4.89 ms. In contrast to this, however, the bound fraction remained unchanged (fig. 9B; 25.3% vs. 23.8% and 24.2%), indicating that cholesterol removal increased the diffusion rate of the probe, without inhibiting its binding. The effect of SM removal also differed strongly for these two probes: TeNT46 in contrast to CTxB, showed significant changes, particularly in surface binding after SMase treatment (18.6% bound; fig. 9B) and in diffusion (τD=6.72 ms). The above results indicate that TeNT46 does not require cholesterol for membrane binding but, unexpectedly, that cholesterol actively slows diffusion of the probe in the membrane. Unlike CTxB, TeNT46 requires SM for effective binding to the membrane. Altogether, the FCS results outlined here are consistent with the lipid requirements identified by the iSPR studies.

### Lipid compositions that favor TeNT46 binding form phase separated domains in GUVs

We wondered whether the pronounced effect of different lipid compositions on the binding of the TeNT46 probe to membranes might have some basis in the propensity of these mixtures to form phase separated domains, since edge-binding preferences have been reported for other toxins (Barlič et al., 2004; Garcia-Saez et al., 2007; Rojko et al., 2014). Therefore, to compare differences in their domain behaviors at longer (i.e. micron) length-scales, Giant Unilamellar Vesicles (GUVs) of ∼5-50 μm diameter (Walde et al., 2010) were prepared with comparable compositions of individual components (POPC/SM/Chol/ganglioside) to those that were used in the iSPR experiments, and imaged in solution (fig. 10). The inclusion of sphingomyelin and ganglioside resulted in a striking phase separation (fig. 10 bottom row), whereas absence of ganglioside or sphingomyelin eradicated visible domains in GUVs, suggesting that peptide binding may be optimized in mixtures that are (at least in artificial systems) able to form sharp domain boundaries.

**Figure 10.**
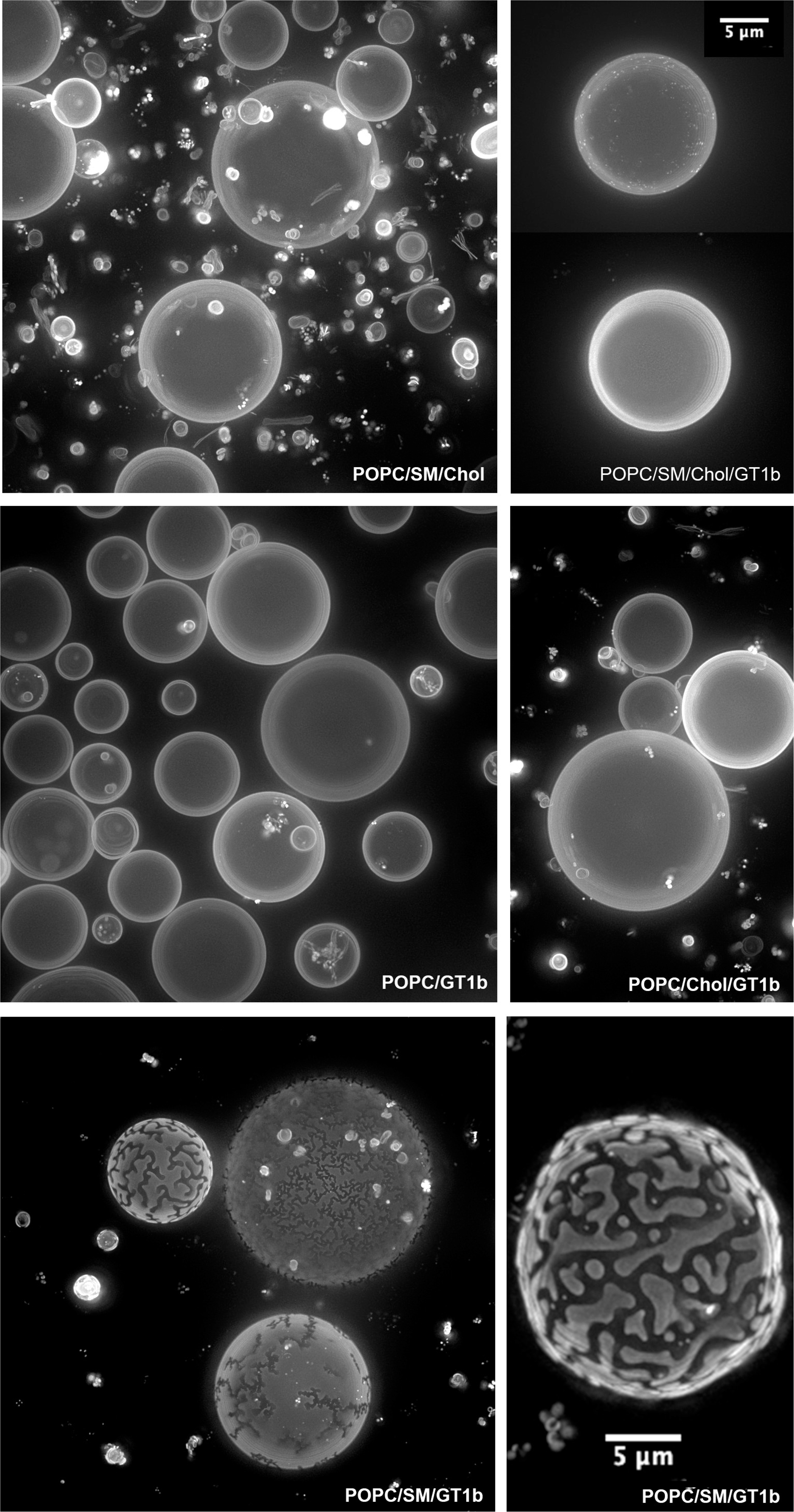
Giant Unilamellar Vesicles prepared with the same mixtures that were used for the iSPR surfaces (see fig. 7). Domain formation at the micron scale is apparent in the POPC/SM/GT1b mixture without cholesterol (bottom row), corresponding to the composition that favoured binding by TeNT46. Domains can be seen clearly in the vesicle in the enlarged image at bottom right.

## Discussion

Lipid domain organization and dynamic features of the plasma membrane of cells are relevant to understanding the mechanisms by which toxins and pathogens, such as the clostridial toxins and ganglioside-interacting viruses, use these interactions to invade cells. We hope ultimately to be able to elucidate differences in toxin-derived probe behaviour and thereby derive a set of rules that dictates diffusion and uptake into the cell, relating these to lipid specificity and binding characteristics. With that long-term aim in mind, we designed a novel probe targeting ganglioside-containing membrane domains, derived from tetanus toxin’s ganglioside-interacting Hc fragment. This probe, designated TeNT46, shows distinctly different binding and diffusion characteristics from the standard GM1 binding microdomain probe CTxB, but importantly retains the specificity of the intact tetanus toxin ganglioside for GT1b.

### The iSPR hybrid bilayer platform

Our characterization of the tetanus-derived TeNT46 probe included the application of an adaptation of SPR ellipsometric imaging or iSPR, combined with a supported hybrid lipid bilayer platform. This was important to be able to assess membrane binding of the TeTN46 toxin-derived probe, and to compare its binding capability and specificity to the known ganglioside binder, CTxB. Critically, the approach applied here was able to reproduce the expected binding characteristics of CTxB, and revealed very different lipid requirements for the new probe, TeNT46. CTxB has a very strong affinity (between 10e-8 M and 10e-9 M) to GM1 ganglioside (Kuziemko et al., 1996), and binds as a pentamer, whereas tetanus has been reported to bind with only ∼10e-7 M to neuronal membranes (Rogers and Snyder, 1981), probably as monomers to GT1b (although some reports suggest that it binds as a dimer (Fotinou et al., 2001) or to two different glycolipids simultaneously (GM1a and GD3 (Chen et al., 2009)). CTxB binding and confined, slow diffusion may require or be enhanced by cholesterol (Bacia et al., 2004; Hebbar et al., 2008; Lingwood et al., 2008; van Zanten et al., 2010), at least within a range of GM1 concentrations (Šachl et al., 2015) but has no requirement for SM, as long as GM1 is present (Stefl et al., 2012). Our method was able to identify these characteristics, supporting its capacity to detect specific toxin-lipid interactions.

Optical biosensors like SPR are well established as an effective means to measure peptide-membrane interactions with supported lipid bilayers (Deleu et al., 2014; Keller et al., 2000; Oliveira et al., 2013). However, SPR, in its traditional form (Patching, 2014) suffers from a number of drawbacks, chief among these being that the integrity of the membrane surface cannot be easily verified, and is in most cases presumed to consist of non-fused vesicles arranged side-by-side (Anderluh et al., 2005; Sofou and Thomas, 2003). Moreover, only a few measurements can be done in one experiment on a single chip, with each binding event representing a stand-alone observation. Another barrier to the study of toxin-membrane interactions in particular is that many of the toxins of interest (e.g. CTxB, Aβ, the clostridial toxins botulinum and tetanus) bind preferentially to charged glycosphingolipids (Kraut et al., 2012; Yowler and Schengrund, 2004), which are difficult to immobilize on standard SPR sensor chips because of their relative hydrophilicity. Here, covalent linking of the lipid-like ODT to the gold chip surface overcame both of these problems, since the SLB surface was both immobile and flat. The <1Å resolution ellipsometric height measurements of the iSPR were able to detect individual layers of the hybrid membrane as it was laid down. Finally, the patches of SLB formed by micro-patterned stamping provided an internal control (the non-stamped areas) for extensive multi-point measurements on different areas of a single chip, greatly improving the significance of the results.

### Lipid requirements of TeNT46 differ from CTxB

The iSPR experiments with different SLB compositions demonstrated a unique lipid specificity of TeNT46, different from that exhibited by CTxB. The lipid targets also differed from those reported for Aβ-and its derivative SBD (Kakio et al., 2002; Steinert et al., 2008). CTxB, as expected, bound strongly to GM1 in our assay, and binding was enhanced by cholesterol, but not significantly by SM. In contrast, TeNT46 bound best to GT1b, like its parent toxin, but absolutely required SM for binding, and in fact was strongly inhibited when SM was replaced by cholesterol in the hybrid bilayer. Surprisingly, the binding preference of the TeNT46 fragment for GT1b containing membranes was the same as that expected for the intact toxin, even though only a subset of the putative ganglioside-contacting residues is included in the 46-residue construct. This may be related to earlier findings (Halpern and Loftus, 1993) that the C-terminal residues of Hc that constitute the trefoil domain (aa 1110-1315) binds to gangliosides more efficiently than a full-length Hc fragment.

### Diffusion and binding measurements by FCS on live cells agree with iSPR binding studies, and distinguish domain dynamics from binding requirements

Using FCS to measure diffusion and binding on live cells, we established that the preferred lipid composition discovered in the hybrid bilayer iSPR assay closely matched cell-membrane (sphingo)lipid/cholesterol requirements revealed by diffusion changes in both CTxB and TeNT46 after lipid perturbations. This is a striking demonstration of the predictive value of the iSPR/hybrid bilayer platform; probe binding to the bilayer reproduced essential lipid compositional features of toxin-cell membrane binding. However, the FCS diffusion data revealed distinct lipid-dependent changes in mobility of the two probes that was independent of binding. CTxB membrane association was expected from earlier studies to depend on cholesterol (Bacia et al., 2004; Hebbar et al., 2008; Lingwood et al., 2008; van Zanten et al., 2010), and indeed this was borne out in both the iSPR and the FCS measurements. In contrast, cholesterol was not required for the binding of TeNT46 in either assay, even though both these peptides interact with gangliosides.

The lack of diminution of TeNT46 binding in response to cholesterol extraction in the FCS experiments was striking, given the expectation that the ganglioside/SM domains would contain cholesterol, according to the raft model (Brown and London, 2000; Simons and Sampaio, 2011; Sonnino et al., 2007), although this is disputed; see (Frisz et al., 2013; Maggio et al., 2004). This finding was especially intriguing, in light of the fact that TeNT46 diffusion, by contrast, was strongly affected by cholesterol depletion. To explain this, we suggest a model whereby binding of the peptide to a specific lipid domain can occur independently of the diffusion of this domain within the plasma membrane. In other words, the platform to which the toxin binds is laterally constrained by lipids (e.g. cholesterol) in the surrounding that do not contribute to binding per se. This lateral restriction could then, in turn, have an effect on other factors like membrane shape fluctuations and clustering dynamics, which influence endocytic uptake by the cell (Johannes, 2017; Pezeshkian et al., 2017).

Interestingly, SM appears to play very different roles in the interactions of the two toxins with the membrane. While not required for CTxB binding or characteristic diffusion by FCS, SM destruction had the greatest effect on TeNT46 binding and diffusion of any of the lipid perturbations. This suggests that a different constellation of lipids constitutes the respective targets of these two probes.

The iSPR/hybrid bilayer methodology was able to accurately reproduce the lipid binding preferences of a well-characterized membrane probe, CTxB, and to define the characteristics of the novel probe, TeNT46. The iSPR platform demonstrated a striking correspondence between the lipid requirements for binding to the artificial SLB membrane, and the binding and diffusion behaviors on actual cells by FCS. Since the iSPR hybrid bilayer platform was able to accurately predict peptide behavior at the cell surface, both methods together (iSPR and FCS) can give information about toxin-membrane interaction that might contribute to—or hinder—cellular invasion by these toxins. We therefore envision this strategy being used to assess the binding affinities and preferences of various toxins, and even to create “super-binding” substrates, in an effort to design decoys.

